# Population dynamics analysis of *Saccharomyces cerevisiae* deletion library during fed-batch cultivation using Bar-seq

**DOI:** 10.1101/2020.04.27.064675

**Authors:** Maren Wehrs, Mitchell G. Thompson, Deepanwita Banerjee, Jan-Philip Prahl, Carolina A. Barcelos, Jadie Moon, Norma M. Morella, Zak Costello, Jay D. Keasling, Patrick M. Shih, Deepti Tanjore, Aindrila Mukhopadhyay

## Abstract

To understand the genetic basis of changes in strain physiology during industrial fermentation, and the corresponding roles these genes play in strain performance, we employed a barcoded yeast deletion library to assess genome-wide strain fitness across a simulated industrial fermentation regime. Our results demonstrate the utility of Bar-seq to assess fermentation associated stresses in yeast populations under industrial conditions. We find that mutant population diversity is maintained through multiple seed trains, enabling for large scale fermentation selective pressures to act upon the community. We identify specific deletion mutants that were enriched in all processes, independent of the cultivation conditions, which include *MCK1, RIM11, MRK1*, and *YGK3* that encode homologues of mammalian glycogen synthase kinase 3 (GSK-3). Further, we show that significant changes in the population diversity during fed-batch cultivations reflect the presence of significant external stresses, such as the accumulation of the fermentative byproduct ethanol. The mutants that were lost during the time of most extreme population selection suggest that specific biological processes may be required to cope with these specific stresses. Overall our work highlights a promising avenue to identify genetic loci and biological stress responses required for fitness under industrial conditions.

## Introduction

*Saccharomyces cerevisiae* is one of the most widely used microbial hosts in biotechnological processes and has been engineered to produce a variety of industrially relevant compounds ranging from pharmaceuticals to biofuels [1–4]. Production strains are typically engineered and optimized in small scale cultivations, while the bioproduction processes take place in large scale bioreactors. Even though it is understood that microbial physiology at larger scales differ from that in shake flask batch cultures, it is typically left to later project stages to optimize the strain and conditions for production at scale [5, 6]. Previous work has indicated that before any candidate strains are employed in industrial scale production environments, it is prudent to adopt de-risking steps that characterize and optimize the strain performance at industrially relevant scales [7]. Given the low throughput and high cost of large-scale bioconversion and fermentation, this step represents a large bottleneck for the development of industrially useful microbial factories [5]. A better understanding of the differences between the culturing conditions in shake flask and large-scale bioreactors will aid our ability to preemptively engineer better microbial hosts before attempting costly and risky scale up.

Multiple functional genomics methods (such as transcriptomics, proteomics, metabolomics, fitness profiling, and fluxomics) have been used to examine critical aspects of strain development such as carbon source utilization, tolerance to toxic substrates or final products, and metabolic flux optimization [8–13]. However, few studies applying omics-level techniques have investigated the differences in process scales on the physiology of a microbial production strain [14–17]. Furthermore, most reports focus on individual aspects of the fermentation process, including oxygen supply [18–20] as well as substrate heterogeneity [21–24]. While these studies shed light on specific physiological changes due to known stresses encountered during the scale up process, such as gas mixing and nutrient heterogeneity, there still remains a dearth of knowledge regarding the biological impact of stresses and bottlenecks on the microbial population at various scales.

Among genome-wide techniques that have proven valuable, massively parallel fitness profiling techniques such as Transposon-Sequencing (Tn-Seq) [25], Barcode-Sequencing (Bar-seq) [26], and Random Barcode Transposon Sequencing (RB-Tn-Seq) [27] allow rapid identification of genetic loci controlling myriad important phenotypes. In this study, we use Bar-seq, which quantifies changes in the population from the changes in abundance of short nucleotide barcodes associated with a known mutation. The Bar-seq methodology has been used extensively since the advent of barcoded deletion and overexpression yeast collections. These collections have led to impressive findings in a wide array of genome-wide phenotypic assays aimed towards increased understanding of biological functions, stress responses, and mechanisms of drug action [28–31]. Similar approaches have been used to assign functions to thousands of genes across many bacteria [32], as well as to identify and eliminate metabolism detrimental to production of a desired molecule [33, 34].

Most industrial biotechnological processes employing microbial production hosts are performed using batch or fed-batch cultivation, which are typically more cost-effective compared to continuous cultivations [35, 36]. In this study, we explored the selective pressures that microbial populations encountered in shake flasks as well as batch and fed-batch bioreactor cultivations to characterize the timing and the identity of these selective pressures. We performed scale-up processes from seed train stage to batch and fed-batch bioreactor cultivation and analyzed the population dynamics of a pooled *S. cerevisiae* deletion collection using Bar-seq. Our results show that the response to these conditions using the yeast Bar-seq library strongly depends on the conditions examined and that certain conditions impose a higher selective pressure.

## Results

We employed the pooled *S. cerevisiae* deletion library to examine the impact of various different cultivation conditions on the physiology of *S. cerevisiae* and compared potential global population differences between cultivations in shake flasks versus bioreactors. We were also able to characterize the impact of process parameters commonly subject to adjustments during fed-batch cultivations on the library. To better simulate industrial processes, we included a two-stage seed train to generate an adequate amount of actively growing cells to inoculate a production bioreactor, as part of our workflow. To avoid any potential impact from amino acid insufficiencies on the fitness of the mutant pools, a prototrophic deletion collection was used in all experiments performed in this study [37].

### Design of Bar-seq Scale Up Fermentation Experiments

It has long been recognized that most strains do not perform the same way when cultivated in a bioreactor compared to a shake flask [6]. The main differences between these two general scenarios are dictated by the geometries of vessels and impellers along with enhanced control capabilities available during cultivations in bioreactors. Better gas mixing [5, 38, 39] can be achieved through presence of impellers and air spargers, better control of nutrient feed rates through external feeding capabilities as well as robust control over pH through the feasibility of acid or base addition. We examined a selection of these conditions between a shake flask cultivation, a bioreactor cultivation in batch mode as well as in fed batch format, and individually across different fed-batch strategies. To assess the effect of any given variable on the population dynamics of a pooled *S. cerevisiae* deletion library, we performed growth competition experiments in two sets, each comprising two seed train stages followed by different final cultivation environments varying in either vessel architecture (set 1: bioreactor versus shake flask) or cultivation parameter (set 2: feeding mode, pH) (Fig 1, for additional details see Fig S1). For each of the two sets, we used an individual aliquot of the pooled library to inoculate the first seed train stage (seed 1) and measured the relative abundances of mutants within the population at least once every 24 h throughout the course of each cultivation using barcode sequencing [40].

**Figure 1:**
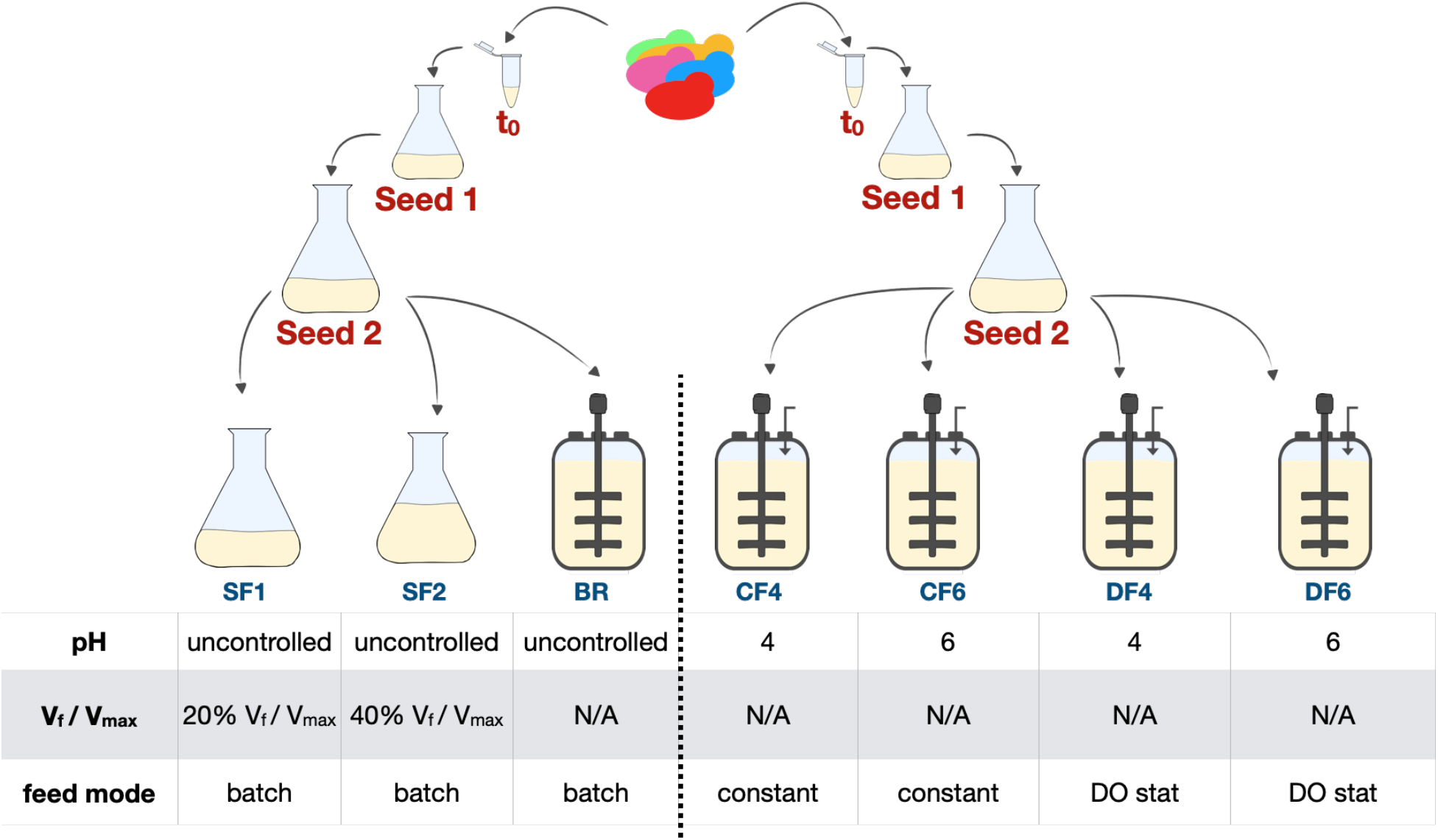
Schematic depiction of experimental setup for the pooled population dynamics experiments. An aliquot of a pooled haploid prototrophic *S. cerevisiae* deletion collection [37] was used to inoculate a two stage shake flask seed train in YPD. Growth competition experiments were performed in two sets, each consisting of two seed train stages (Seed 1 and Seed 2) followed by different main cultivation environments. Set 1 (left) varied in vessel architecture that included shake flasks with different culture volumes (SF1 and SF2) and a batch bioreactor (BR). Set 2 (right) varied in cultivation parameter settings that included four fed-batch mode bioreactors (CF4, CF6, DF4 and DF6) with two different feeding modes (CF = constant feed, DF = DO- signal based feed) and two different pH (4 and 6). All cultures were run at 30°C, the dissolved oxygen (DO) was not controlled. Samples were taken at the end of each seed stage and throughout the bioreactor runs.

In contrast to microbial cultivation in shake flasks, where gas transfer is achieved predominantly by shaking, bioreactors can employ oxygen spargers and agitators to ensure improved oxygen transfer and uptake by the cultures. To test if these differences in gas mixing impacts the population dynamics of the mutant pool, we performed batch cultivations, with no DO control, even in the bioreactors. A single bolus of substrate was added to the culture media at the beginning of the cultivation and the composition of the resulting mutant pool after 72 h was compared. Temperature was the only parameter controlled in the bioreactor, leaving pH uncontrolled as well. To administer differences in oxygen transfer rates, we tested two different shake flask filling volumes (20 % V_f_ / V_max_ and 40 % V_f_ / V_max_).

Bioreactors also allow for pH control to ensure an optimal production environment as well as external feed of additional substrate to extend the process by minimizing nutrient limitations. To assess the impact of a feeding regime and pH on the population dynamics of the mutant pool, we performed four fed-batch cultivations: (i) two pH set points, pH 4 and pH 6, and (ii) two glucose feeding schemes, constant rate feeding and a DO signal-based pulse feeding (Fig 1, Fig S1 and Fig S3). In a DO signal feeding regimen, a feeding solution is added upon complete exhaustion of available carbon sources and thus is responsive to the metabolic activity of the cells. Whereas, under constant rate feeding regimen, feed solution is continuously added to the fermentation broth independent of the metabolic activity. If the feed rate is not optimized for the process, then this feeding strategy tends to result in large accumulation of fermentative by-products, including ethanol, due to overfeeding of the culture. In all cases, the fed-batch phase was triggered after initial DO spikes, indicating full consumption of the carbon sources available during the batch phase.

### Mutant Population Succession is Dictated by Fermentation Regime

We interrogated the changes detected in the mutant population structure of each cultivation over time and visualized the data using a correlation matrix showing the diversity of the mutant pool via the total number of observed mutants. Of the 6002 total barcoded genes present in this deletion collection, 3099 were detected at t0 with at least 10 counts. To minimize statistical noise, we only included genes with a threshold of at least 10 counts in the Seed 2 population in this analysis. It may be noted that the general trend of observed mutants per condition is robust and independent of this threshold (Fig S4).

In the studies where vessel geometry was the only variable, microbial growth, measured by optical density (OD), and glucose consumption profiles look similar for all batch cultivations, independent of the cultivation vessel (BR vs. SF1 and SF2 in Figure S2a). Overall, the population structure of the two individual seed trains were highly similar (r > 0.99 across all seed populations) when compared via Pearson correlation of mutant barcode counts (Fig S5, Table S1). The overall structure of the populations cultivated in both shake flask conditions (SF1 and SF2) and the batch bioreactor (BR) were also highly similar when compared via Pearson correlation (Fig 3) of mutant barcode counts (r > 0.99 across all time points), with minor changes at 48 h (for additional details see Fig S2b and c). As the overall population structure is highly similar in all conditions and time points tested, we conclude that the difference in vessel architecture is not reflected in the population structure of the deletion mutant pool.

For population changes between batch, and a fed- batch environment for the tested conditions, we analyzed the abundance of individual barcode counts in populations under the respective conditions, comparing BR to CF 4, 6 and DF 4, 6. Overall, only 8.6 % (212 barcodes) of mutant barcodes initially present in the respective seed 2 culture were retained in all populations cultivated in a bioreactor environment independent of the cultivation mode (batch versus fed-batch), and 0.53 % (13 barcodes) of mutant barcodes were lost in all conditions at the final timepoints (Fig 2B). Upon examination of the mutant populations in the fed-batch conditions specifically, we observed a decrease in the overall diversity of our mutant population over the course of the fermentation for all four conditions (Fig 2B). As expected from findings from the batch bioreactor study (BR) and the correlation matrix (Fig 3, Fig. S2b), over 90% of the mutants in the initial Seed 2 mutant population are still present throughout the batch phase (< 33 h EFT) with a highly similar population structure across all conditions (CF 4, 6 and DF 4, 6) before feeding began. Even then, all samples taken up to 48 h EFT (elapsed fermentation time) show high correlation to each other, indicating little change in the pool of detected barcodes. However, large differences in population composition were observed in the populations grown in a constant rate feeding regime, in CF4 and 6, by the end of the fermentation at 119 h. The constant rate feed resulted in four times higher glucose addition (~250 g) but only slightly higher OD values (OD_600_ ~ 52) than DO signal based feeding (OD_600_ ~ 40, < 75 g glucose added). Most of the glucose in CF 1 and 2 accumulated as the fermentative by-product ethanol (~37g/L), much higher compared to that in DF4 and 6 (< 1g/L ethanol produced) (Fig 2A, Fig S1). It is possible that the presence of high concentrations of ethanol in CF 1 and 2 led to the loss in diversity.

**Figure 2:**
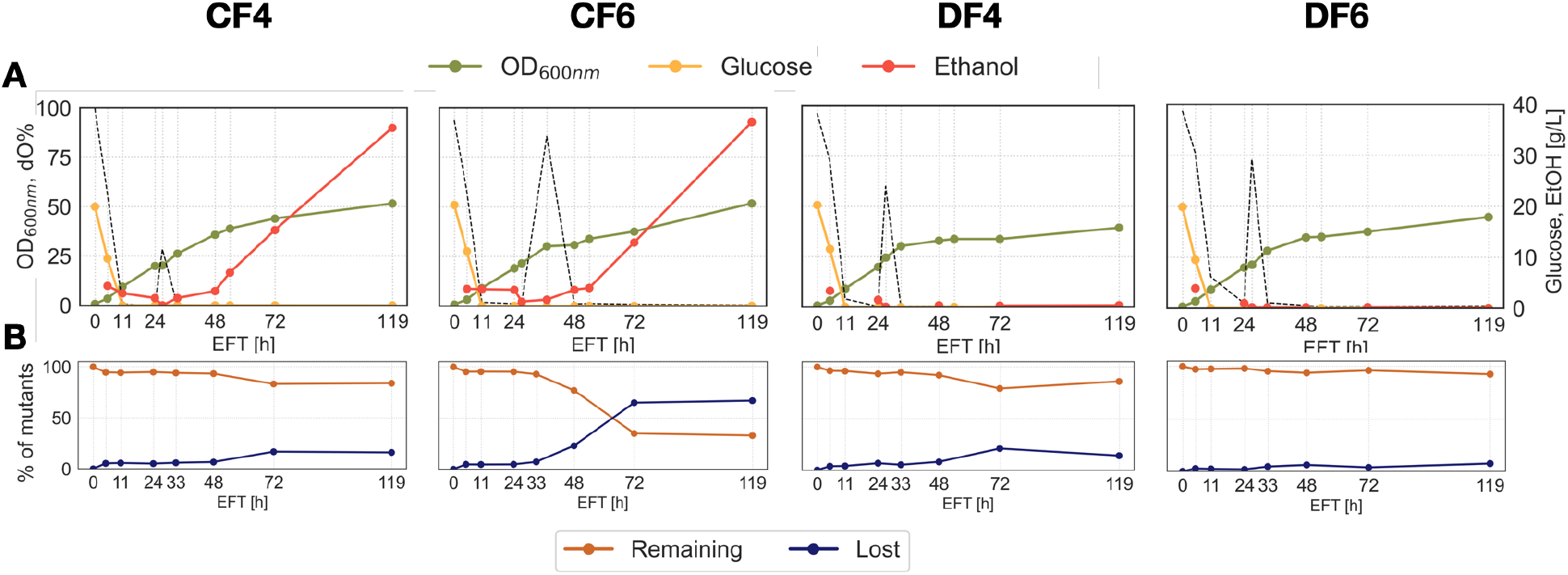
A: Growth profiles of each fed-batch bioreactor over time. Concentrations of glucose (yellow line) and ethanol (red line), OD_600_ (green line) and the culture %DO (dotted line) are plotted against time for cells grown in different fed-batch regimes. **B: Population diversity of each fed-batch bioreactor over time.** Depicted are the counts for barcodes that were detected (blue) and not detected (orange) at a given time point at a threshold of n = 10.

**Figure 3:**
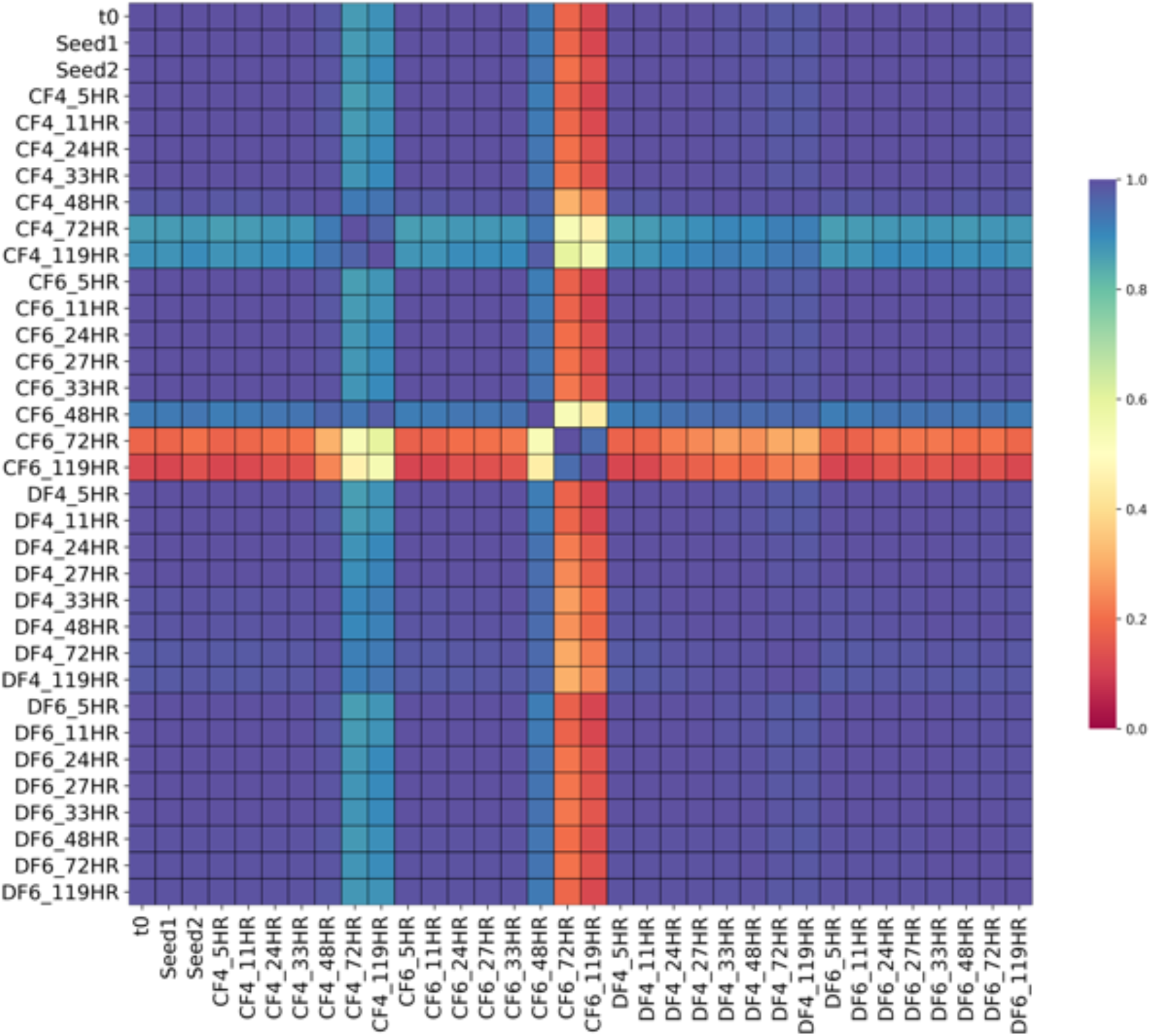
Correlation matrix of mutant abundance. Heatmap shows pairwise Pearson correlation coefficients of the raw number of mutant barcodes counted in batch and fed-batch cultures tested with constant rate and DO signal feeding at two levels of pH, 4 and 6. Any gene that had less than 10 barcode counts in every condition was not considered for analysis. Scale bar shows Pearson correlation coefficient from 1 to 0.

Both populations grown under a DO signal-based feeding regime, DF4 and 6, remained highly correlated to the initial population detected in Seed 2 and each other throughout the cultivation independent of the culture pH (DF4 = pH 4, DF6 = pH 6, r = 0.98). Further, less than 200 mutants (i.e. less than 7.5% of the mutants present in the Seed 2 population) in reactor DF6 were lost over the course of the entire cultivation, indicating that the conditions (pH 6, DO signal based feeding) did not result in sufficient selective pressure for a significant change of the mutant pool population over the course of 119 h. In both reactors cultivated at pH 4, CF4 and DF4, an additional 10% of the mutants in the respective mutant population were lost within a 24 h window in the fed-batch phase between 48 h and 72 h compared to DF6. In contrast, the population cultivated in CF6 (pH 6, constant rate feed) exhibited the start of diversity loss earlier, losing 23% of initial mutant barcodes between 33 h EFT and 48 h EFT and additional 55% of these mutants between 48 h and 72 h, resulting in a decrease of its diversity to about 35% of its initial mutant pool after 72 h. This observation is also reflected in the correlation matrix: the mutant population of CF4 (pH 4) showed a modest decrease in correlation to DF6 after 48 h (pH 6, *r* = 0.88), whereas the CF6 population (pH 6) changed dramatically in comparison to DF6 at that time point (*r* = 0.12), and only showed moderate correlation to CF4 (r = 0.54). These observations indicate that, for this yeast deletion collection, the selection of the feeding scheme that leads to the presence of high concentrations of the byproduct – ethanol impacts the diversity of the mutant pool more than the pH of the culture broth.

Upon comparison of the barcode abundance trends present under the conditions in CF4 and 6, we found that the majority of loss in diversity takes place between 48 h and 72 h for both of these populations, which is representative of the onset of stationary phase. This time point seems to coincide with significant accumulation of ethanol in these 2 bioreactor environments reaching ethanol concentration of over 12 g/L by 72 h from 3g/L at 48 h in both reactors (Fig 2A, red line).

### Deletion mutants of genes associated with mitochondria are commonly lost across all conditions tested

Amongst the deletion mutants that were maintained throughout the entire fermentation, we observed that each condition (shake flask, batch bioreactor as well as fed-batch processes) resulted in overlapping as well as distinct mutant pools with increased or decreased fitness (Fig 4). Our analysis revealed that 47 mutants were overrepresented specifically in the batch bioreactor, 4 mutants in the shake flask and 30 mutants were uniquely overrepresented in the fed-batch conditions at the final timepoint. While almost 50% of the mutants overrepresented in the shake flask population, were also overrepresented in the batch bioreactor population (3 of 7 barcodes), no overlap was detected for overrepresented mutants between the fed-batch bioreactors and the shake flask. However, out of 47 mutants overrepresented in the fed-batch environment, 17 were also overrepresented in the batch bioreactor but not in shake flasks.

**Figure 4:**
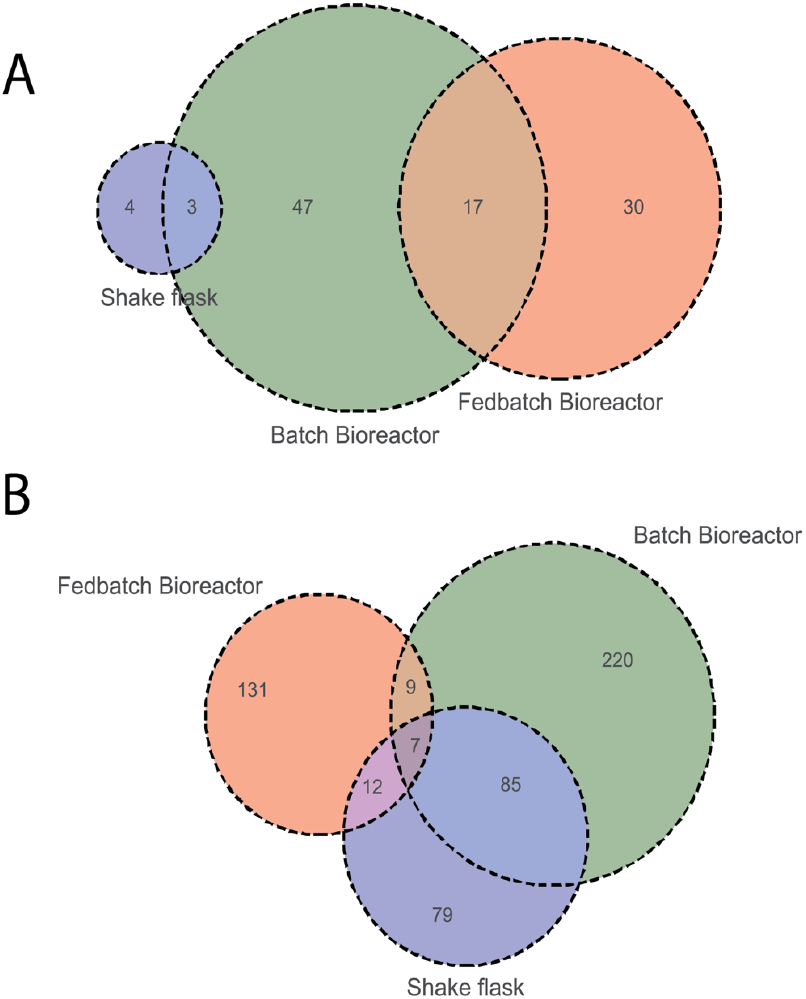
Extant mutants that were commonly more (A) or less (B) fit over the course of the cultivation in either fed-batch bioreactors, batch bioreactor, or shake flasks from 0 to 119 h for the fed-batch bioreactors and 0 h to 72 h for the batch bioreactor and shake flasks. As shown in Figure 2B, the majority of population changes occurred before 72 h in fed-batch cultivations. Venn diagrams show the number of mutants common or unique between parameters tested.

To better understand the biological relevance of these commonly lost and commonly retained mutants, we performed gene ontology enrichment analysis of these gene lists using the Benjamini-Hochberg procedure to decrease the false discovery rate of our analysis. We found that only 3% of the barcodes initially present in Seed 2 (n= 77) were commonly lost across all the fed-batch bioreactors while 31% of the barcodes (n = 796) were commonly retained by the end of the experiment (119 h) (Fig 2B). Of the 77 deletion mutants commonly lost across all fed-batch bioreactors, almost 50% (28 genes) are annotated to be associated with mitochondria. Of these 28, the majority of genes (20) is annotated to associate to the mitochondrial envelope and 9 genes are associated with mitochondrial organization including the assembly of the mitochondrial respiratory complex. Of the 796 deletion mutants retained in all fed-batch bioreactors with 787 unique gene identifiers, the annotated genes associated with response to salt stress as well as diphosphate activity were observed as significantly enriched.

Amongst the 10 genes annotated to be associated to cellular response to salt stress (GO ID: 0071472) are *MSN4*, a stress-responsive transcriptional activator involved in the yeast general stress response as well as all four genes *MCK1, RIM11, MRK1*, and *YGK3*, that all encode homologues of mammalian glycogen synthase kinase 3 (GSK-3). Yeast GSK-3 homologues are suggested to promote formation of a complex between the stress-responsive transcriptional activator Msn2p, a paralog to Msn4p and DNA, which is required for the proper response to different forms of stress [41, 42]. These five deletion mutants (*msn4*^−^, *mck1*^−^, *rim11*^−^, *mrk1*^−^, *ygk3*^−^) were also retained throughout the study performed in the batch bioreactor and SF2, making these genes general beneficial targets for gene deletion to improve fitness independent of glucose availability, pH or vessel architecture.

### Bioreactor-specific Stress Responses Revealed by Bar-seq

In reactors CF4 and 6, which were run under a constant rate feeding scheme, the drastic loss of diversity coincided with the onset of ethanol accumulation. In DF4, the implementation of a DO signal feeding scheme resulted in no substantial accumulation of ethanol but rather a frequent switch from respiratory to fermentative metabolic state triggered by the addition of small glucose boluses that are characteristic of a DO signal feeding scheme (Fig S3). To determine whether the groups of mutants that were either lost or selected against during this cultivation period in the respective populations are indicative of the specific stresses imposed on the microbial population, we performed a direct comparison of the mutants specifically reduced or lost in each of the bioreactors followed by gene ontology enrichment analysis of those mutant groups. To allow the determination of statistically significant enrichments of genes in these groups, we excluded the population in bioreactor CF6 from further analysis, as it was reduced to less than 50% of its initial mutant diversity by 72 h.

Overall, 174 mutants were uniquely selected against in bioreactor CF4, 253 in bioreactor DF4 and only 39 mutants were either lost or appreciably selected against in the population cultivated in bioreactor DF6 (Fig 5). Our analysis revealed that 112 mutants were selected against in both populations cultivated at pH 4 (CF4 and DF4), while only 2 mutants were commonly lost or reduced in all bioreactors at this time period. Glucose addition in DF6 was controlled by measurements of dissolved oxygen within the culture broth (DO signal), which is an indirect indicator for metabolic activity of the cells in suspension. The frequency of DO spikes, and thus the rate at which the cells are consuming all the carbon delivered per feed bolus, significantly decreases over time in case of DF6 compared to DF4, starting at around 50 h EFT. The observed low number of mutants lost in bioreactor DF6 could be attributed to the comparably lower metabolic activity of that culture (Fig S3).

**Figure 5:**
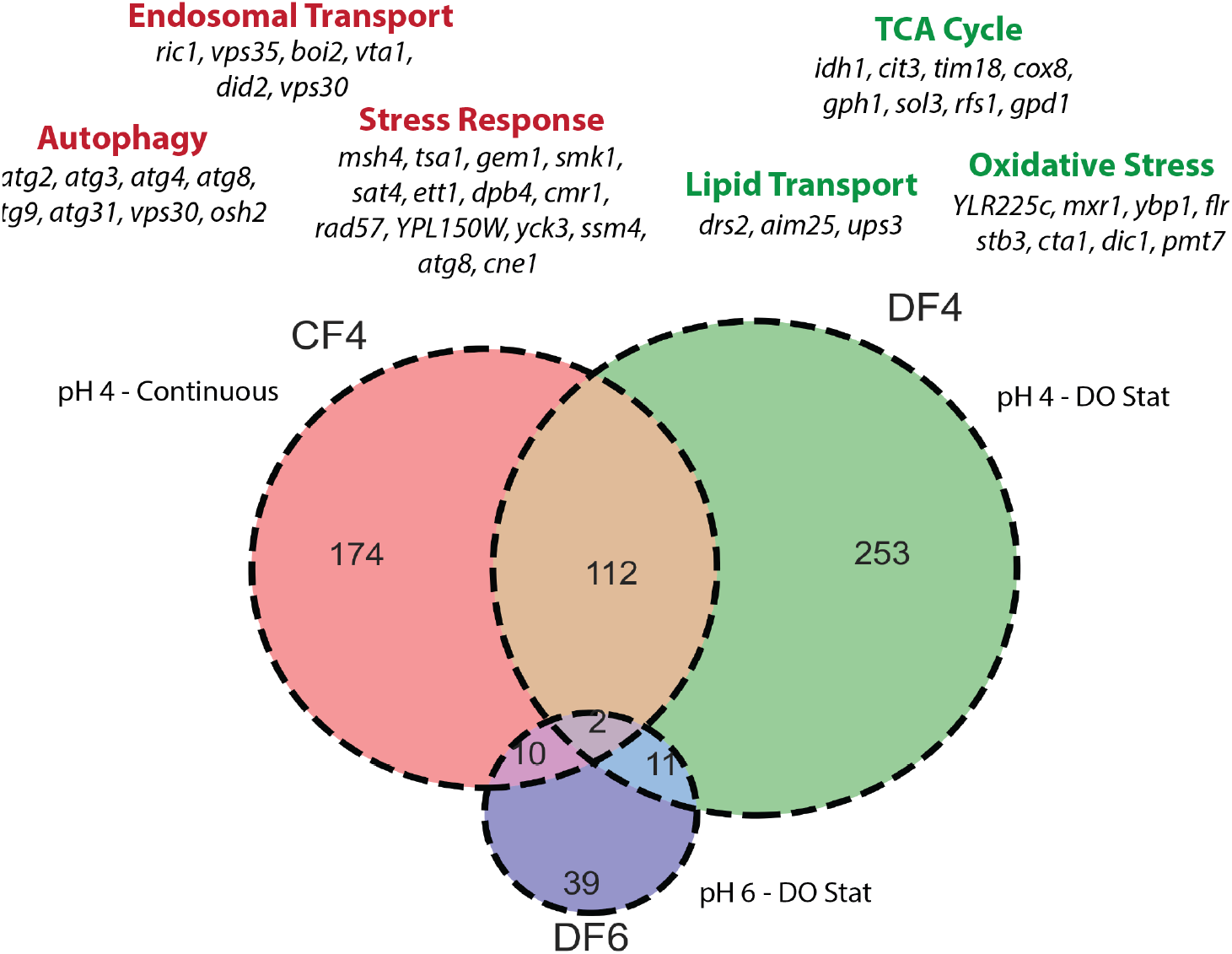
Mutants selected against in bioreactors from 48 h to 72 h. Venn diagram shows mutants that either were lost (had a total barcode count <10) or were 50% less relatively abundant from 48 h to 72 h. Red text shows GO terms that were enriched in lost or less abundant mutants in bioreactor CF4, while green test shows enriched GO terms from bioreactor DF4. Specific genes from each GO term are written below in black.

The culture in bioreactor CF4 was maintained at pH 4 and was fed at a constant rate with glucose, which resulted in a high accumulation of the fermentative by-product ethanol starting at 48 h. The group of mutants selected against during this period of cultivation in CF4 is enriched with genes associated with cellular stress (5.14-fold enrichment, *p* = 0.0415), autophagy (5.9-fold enrichment, *p* = 0.0023), and endosomal transport (6.66-fold enrichment, *p* = 0.00144). Of those genes associated with cellular stress, the majority is either associated with DNA damage (*MSH4, TSA1, GEM1, DPB4, CMR1, RAD57*) or protein misfolding (*CMR1, SSM4, CNE1*). Additionally, we observed an overrepresentation of genes involved in the Cytoplasm-to-vacuole targeting (Cvt) pathway, a constitutive and specific form of autophagy that uses autophagosomal-like vesicles for selective transport of hydrolases to the vacuole, in the pool of mutants that were specifically selected against in bioreactor CF4 between 48 h and 72 h *(ATG2, ATG3, ATG4, ATG8, ATG9. ATG31, VSP30)*. Similar to bioreactor CF4, the culture in bioreactor DF4 was also maintained at pH 4 but was fed intermittently based on DO spike signals. Here, the pool of mutants that were significantly selected against was enriched for genes associated with the TCA cycle (6.09-fold enrichment, *p* = 0.0182) as well as oxidative stress (4.68-fold enrichment, *p* = 0.0142). The group of mutants selected against predominantly comprised of genes associated with the mitochondria (*CIT3, IDH1, COX8, TIM18, UPS3, AIM25, DIC1*) as well as general stress response genes (*GPH1, GPD1*) and oxidative stress responses (*MXR1, YBP1, CTA1*).

In addition, we found that 220 mutants were underrepresented specifically in the batch bioreactor, 79 in the shake flask and 131 were uniquely underrepresented in the fed-batch conditions at the final timepoint (**Fig 4**). Unlike the overrepresented mutant pool analysis, a subset of underrepresented mutants is shared in two conditions and 7 mutants are underrepresented in all three conditions. The majority of these mutants, namely rps0A^−^, rpl8B^−^, esa1^−^, ric1^−^, msc6^−^, can be associated with ribosomal and translational activity [43–46]. Similar to the overrepresented mutant pool at the final time point, we observed a large overlap of the underrepresented mutant pool in the shake flask population and batch bioreactor population (47%, 85 of 183 underrepresented in shake flasks). The pool of 131 mutants, specifically underrepresented but still present in fed-batch bioreactors, was enriched with genes associated with mitochondria (*p*=0.01, n=39, GO ID = 0005739). Of the 39 mitochondria associated genes, 11 play a role in mitochondrial organization while 7 are associated with cellular respiration

## Discussion

In this study we demonstrate the feasibility to correlate physiological changes observed under controlled cultivation conditions, including different pH set points and fed-batch regimes, with significant changes in population structures of yeast barcoded mutants. Specifically, we found that the choice of feeding strategy generating toxic by-products had a great effect on the population structure, highlighting the importance of feed rate adjustments and dynamic feeding strategies during the biocatalysis phase in fed-batch processes.

We found the *S. cerevisiae* population structure to be robust throughout the entire seed train as indicated by a Pearson correlation of population makeup (Fig 3). This finding is critical as significant perturbations in the population structure at seed train stage would render the analysis of subsequent scale-up experiments infeasible. Interestingly, while no significant change in population structure was observed in the seed train, changes in the mutant pool composition were observed in all conditions mimicking production cultivations, including shake flask and bioreactor experiments, allowing us to identify mutants in genes that were specifically selected against in each of the tested conditions.

We found that the pool of deletion mutants specifically selected against in fed-batch conditions tested in this study was enriched with genes associated with mitochondria, specifically in mitochondrial organization and cellular respiration. As a Crabtree-positive organism, *S. cerevisiae* predominantly metabolizes glucose by fermentation even under aerobic conditions [47], which causes the cells to be in a non-respiratory metabolic state during typical batch fermentation processes that employ glucose as a carbon source [48]. We observed that deletion mutants associated with the mitochondria are specifically selected against in fed-batch regimes and not in the batch bioreactors. Generally, we did not observe significant changes in the population in any tested environment until glucose depletion occurred, which constitutes the initiation of the feeding phase in fed-batch processes (Fig 2, Fig S3). The apparent selection against mutants harboring deletions of genes associated with mitochondrial organization and cellular respiration in fed-batch processes could be an artifact of the extended growth phase caused by the addition of glucose in the fed-batch processes: While we did not detect an enrichment of genes associated in the first “wave” of genes selected again, there might be a “second round of losers” which only became apparent due to the additional generations. As mitochondria harbor important metabolic pathways for various native and non-native bioproducts [49–51], better understanding of the cause of this negative selection is required to enable better process and strain designs aimed at maintaining functional mitochondria during scale up.

An important metric to determine the suitability of Bar-seq to elucidate stresses involved in industrial scale-up was that loss of population diversity correlated to the onset of observable stress. Temporal analyses of mutant populations showed a greater than 10% loss of overall mutant diversity in 3 out of 4 fed-batch bioreactors from 48 to 72 h of fermentation runs (Fig 2). In the constant rate feed bioreactors CF4 and 6, this observation correlated with the onset of ethanol production, which may have imposed a strong selective pressure against many mutants. Our results suggest that DO signal feeding is a far less selective environment than constant rate fed-batch fermentations in which DO was not controlled.

We detected that deletion mutants of genes associated with autophagy and a general stress response were specifically selected against under constant rate feeding conditions (CF4, Fig 5). Autophagy is the process whereby cytoplasmic components and excess organelles are degraded and is known to be initiated upon starvation for nutrients such as carbon, nitrogen, sulfur, and various amino acids, or upon endoplasmic reticulum stress [52, 53]. Our findings are in agreement with Piggott *et al,* who found genes that function in autophagy to be required for optimal survival during fermentation when performing a genome-wide study of *S. cerevisiae* gene requirements during grape juice fermentation [54]. The authors concluded that the recycling of cellular components by autophagy enables yeast to survive the stressful conditions of fermentation and maximize fermentative output [54].

Our work was able to identify genetic loci that are selected for or against in various bioreactor schemes. However, the time and cost associated with each individual fermentation run precluded biological replicates and established stochastic variability occurring over prolonged bioreactor studies. While our study confirms earlier findings that genes involved in autophagy can be associated with increased fitness during fermentation, future work should focus on robust replication to increase statistical relevance and allow identification of deletion mutants with more subtle impact on fitness in an industrial setting.

## Materials and Methods

### Yeast Strain Collections

The prototrophic yeast deletion collection in the haploid *S. cerevisiae* BY4742 background was obtained as individual colonies from the Caudy lab (Univ of Toronto) plated on Omnitrays containing YPD agar with 200 mg/mL G418. Balanced pools of the deletion collection were obtained by harvesting all individual colonies into a total of 100ml YPD containing 200 mg/mL G418 resulting in a cell suspension of OD_600_ = 50. 1 ml aliquots of the yeast deletion pools with a final concentration of 25 % glycerol were prepared and stored at −80 °C until further use.

### Seed cultivation

The yeast strain collection was cultured in a two-tiered seed train. The first seed culture was grown in 250 mL baffled shake flasks containing 50 mL YPD media (10 g/L yeast extract, 20 g/L peptone and 20 g/L glucose) inoculated with a 0.7 % (v/v) inoculum directly from the glycerol stock. The second seed culture was grown in 500 mL baffled shake flasks containing 100 mL YPD media using a 10 % (v/v) inoculum size. All seed cultures were incubated at 30 °C, 200 rpm (1” throw) for 24 h.

### Shake flask experiments

Shake flask experiments were carried out in 500 mL baffled Erlenmeyer flasks containing 100 mL (V_f_/V_max_ = 20 %) and 200 mL (V_f_/V_max_ = 40 %) of YPD media with an initial pH of 6.1. V_f_ is defined as the volume of media in the flask and V_max_ is the total volume of the flask. Flasks were inoculated from second seed cultures (4 % (v/v) and incubated at 30 °C and 200 rpm (Orbit diameter 25 mm). Cultures for serial dilution experiments were grown for 24 h and back diluted into a flask with fresh YPD media. This was done two times until 72 h total fermentation time was reached. Samples were taken once a day and centrifuged at 15,000 x g for 5 min. Supernatant was stored at 4 °C and the cell pellet was stored at −80°C.

### Bioreactor experiments

Batch and fed-batch bioreactor experiments were performed in 2 L bench top glass fermentors (Biostat B, Sartorius Stedim, Göttingen, Germany) equipped with two 6-blade Rushton impellers. All tanks were batched with 960 mL of YPD media and autoclaved at 121°C for 30 min. The bioreactors were inoculated with second seed cultures with inoculum size of 4 % (v/v) initial batch volume, 40 mL in 1 L. Temperature, agitation and air flow were maintained constant at 30 °C, 400 rpm and 0.5 vvm, respectively and pH was controlled to 4.0 and 6.0 using 2 N NaOH and 2 N HCl. The batch bioreactor experiment started with little less than 20 g/L sugar, which was consumed within the first 20 h of the study. However, ethanol and other carbon sources generated in the first few hours lasted for up to 72 h, with DO not reaching 100% until then. To mimic the shake flask studies, for BR, glucose was not replenished.

Glucose was replenished in fed-batch experiments: CF1, CF2, DF1, and DF2. Two different feeding strategies, i.e. a linear feed profile (y = 0.117 mL/h^2^ x + 4 mL/h) and a dissolved oxygen (DO) signal-triggered pulsed feeding loop (ΔDO = 30 %, Flow rate = 40mL/h, Pump duration = 6 min). Linear feed was conducted at a constant rate that was chosen based on previous experience to avoid glucose starvation of the microbial culture. A DO signal based pulsed feeding can be explained as follows: upon all carbon (glucose, ethanol, organic acids, etc.) in the culture is consumed, the metabolic activity stalls. Since the reactor is continuously pumped with air, no oxygen consumption due to stalled metabolic activity leads to a sharp increase in dissolved oxygen signal. An automated system ensures that once the increase exceeds a certain threshold value, the feed controller is signaled to administer a predetermined bolus of substrate. This type of DO signal based pulsed feeding continues until the fermentation process is ended. The initial DO spikes for CF2, DF1, and DF3 occurred within a 2-hour time period. The DO spike for CF1, however, occurred 8 h later due to the presence of excess ethanol in the media. The feed media composition was as follows: 10 g/L yeast extract, 20 g/L peptone and 250 g/L glucose. 1 ml samples were taken in regular intervals (5 h, 24 h, 33 h, 48 h, 72 h and 119 h) and centrifuged at 15,000 x g for 5 min. The supernatant was filtered (0.2 mm) and stored at 4°C. The cell pellet was stored at −80 °C.

### Preparation of genomic DNA and Bar-seq analysis

Genomic DNA was extracted from dry, frozen cell pellets using the “Smash- and-Grab” method published by Hoffman and Winston [55] and used for barcode verification. Amplifications of the barcodes using previously published primers [31] and sequencing of the libraries using an Illumina MiSeq was performed at the Vincent J. Coates Genomics Sequencing Laboratory, California Institute for Quantitative Biosciences (QB3) (Berkeley, California, USA). Each barcode was reassigned to a gene using a standard binary search program as described by Payen *et al.* [31]. Only reads that matched perfectly to the reannotated yeast deletion collection were used [25]. Multiple genes with the same barcodes were discarded. Strains with less than 10 counts in the starting pool (t0) were discarded. The numbers of strains identified in the conditions are summarized in Table S1. To avoid division by zero errors, each barcode count was increased by 10 before being normalized to the total number of reads for each sample.

### Statistical analysis

Pearson correlations were calculated with the SciPy Python library [56]. To identify the time points of “mass extinction” / biggest steps in loss of diversity (here, defined as gene richness), we looked at the number of genes either lost or retained during the time course for each of the bioreactors using different count thresholds for mutants present in the Seed train (>0, 5, 10 or 20).

### Gene Ontology enrichment

Gene ontology enrichment analysis was performed using the List analysis tool provided by YeastMine (http://yeastmine.yeastgenome.org), populated by SGD and powered by InterMine [57] against the annotated *S. cerevisiae* S288C genome. The tests were corrected using the Benjamini-Hochberg procedure to decrease the false discovery rate of our analysis with a maximum acceptable p-value of 0.05 unless indicated otherwise.

## Acknowledgements

We would like to thank Amy Caudy (University of Toronto) for generously sharing the auxotrophic yeast deletion collection that was used in this study as well as Shana McDewitt for helpful discussions and library preparations and subsequent sequencing. We thank Robert Haushalter for critically reviewing this manuscript. This work was part of the DOE Joint BioEnergy Institute (https://www.jbei.org) supported by the U.S. Department of Energy, Office of Science, Office of Biological and Environmental Research, and was part of the Agile BioFoundry (http://agilebiofoundry.org) supported by the U.S. Department of Energy, Energy Efficiency and Renewable Energy (EERE), Bioenergy Technologies Office (BETO), through contract DE-AC02-05CH11231 between Lawrence Berkeley National Laboratory and the U.S. Department of Energy. Bioreactor and shake flask studies were conducted at the Advanced Biofuels and Bioproducts Demonstration Unit (ABPDU). EERE BETO also supports ABPDU operations. The views and opinions of the authors expressed herein do not necessarily state or reflect those of the United States Government or any agency thereof. Neither the United States Government nor any agency thereof, nor any of their employees, makes any warranty, expressed or implied, or assumes any legal liability or responsibility for the accuracy, completeness, or usefulness of any information, apparatus, product, or process disclosed, or represents that its use would not infringe privately owned rights.

## Contributions

M.W., J.P.P. and D.T. designed the experiments, M.W., J.P.P., C.A.B. and J.M. performed the experiments, M.W., M.G.T, D.B., Z.C. and N.M. analyzed data. M.W., M.G.T and AM composed the manuscript. P.M.S., J.D.K, D.T. and A.M. guided the scope of the project, provided critical input for the manuscript and provided resources. All authors read and approved the final manuscript.

