## Supplementary Info for "Population dynamics analysis of *Saccharomyces cerevisiae* deletion library during fed-batch cultivation using Bar-seq"

[1]Biological Systems and Engineering Division, Lawrence Berkeley National Laboratory, Berkeley, CA 94720, [2]Joint BioEnergy Institute, Lawrence Berkeley National Laboratory, Emeryville, CA 94608, [3]Advanced Biofuels and Bioproducts Process Development Unit, Lawrence Berkeley National Laboratory, Emeryville, CA 94608, [4], Fred Hutchinson Cancer Research Center, Seattle, WA 98109, USA [5]Department of Energy Agile BioFoundry, Emeryville, CA 94608, [6]Department of Plant and Microbial Biology, University of California, Berkeley, CA 94720, USA, [7]Department of Bioengineering, University of California, Berkeley, CA 94720, USA, [8]Department of Chemical and Biomolecular Engineering, University of California, Berkeley, CA 94720, USA, [9]The Novo Nordisk Foundation Center for Biosustainability, Technical University of Denmark, Denmark, [10]Synthetic Biochemistry Center, Institute for Synthetic Biology, Shenzhen Institutes for Advanced Technologies, Shenzhen, China [11]Environmental Genomics and Systems Biology Division, Lawrence Berkeley National Laboratory, Berkeley, CA 94720, [12]Department of Plant Biology, University of California-Davis, Davis, CA 95616, USA

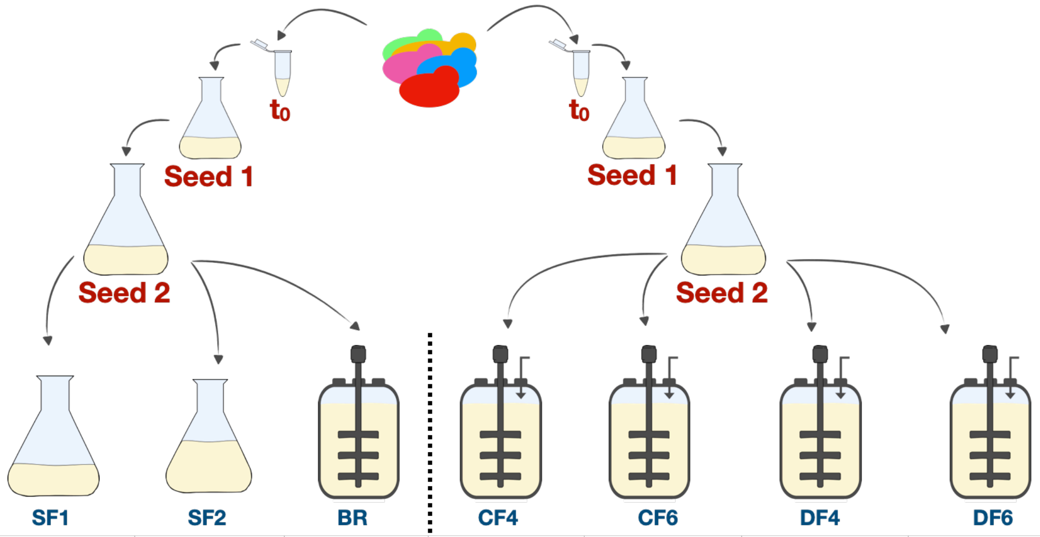


| | SF1 | SF2 | BR | CF4 | CF6 | DF4 | DF6 |
|---|---|---|---|---|---|---|---|
| **pH** | uncontrolled | uncontrolled | uncontrolled | 4 | 6 | 4 | 6 |
| **$V_f$ / $V_{max}$** | 20% $V_f$ / $V_{max}$ | 40% $V_f$ / $V_{max}$ | N/A | N/A | N/A | N/A | N/A |
| **feed mode** | batch | batch | batch | constant | constant | DO stat | DO stat |
| **$EFT_{total}$** | 72 h | 72 h | 72 h | 119 h | 119 h | 119 h | 119 h |
| **$OD_{600,\ final}$** | 21.9 | 23.7 | 21.4 | 51.8 | 51.6 | 39.4 | 44.8 |
| **Glucose $_{added}$** | 2 g | 4 g | 20 g | 243 g | 250 g | 74 g | 69 g |
| **Ethanol $_{final}$** | N/A | N/A | N/A | 37.1 g/L | 35.9 g/L | 0.3 g/L | 0.01 g/L |
| **Acetic Acid $_{final}$** | N/A | N/A | N/A | 0.7 g/L | 2. 1 g/L | 0.6 g/L | 0.6 g/L |

**Figure S1: Overview of all conditions tested in this study.** Growth competition experiments were done in two sets, each consisting of two seed train stages (Seed 1 and Seed 2) followed by different main cultivation environments. Set 1 (left) varied in vessel architecture that included batch bioreactor (BR) and shake flasks with different culture volumes (SF1 and SF2). Set 2 (right) varied in cultivation parameter setting that included four fed-batch mode bioreactors (A05, A06, A07 and A08) with two different feeding modes and two different pH.

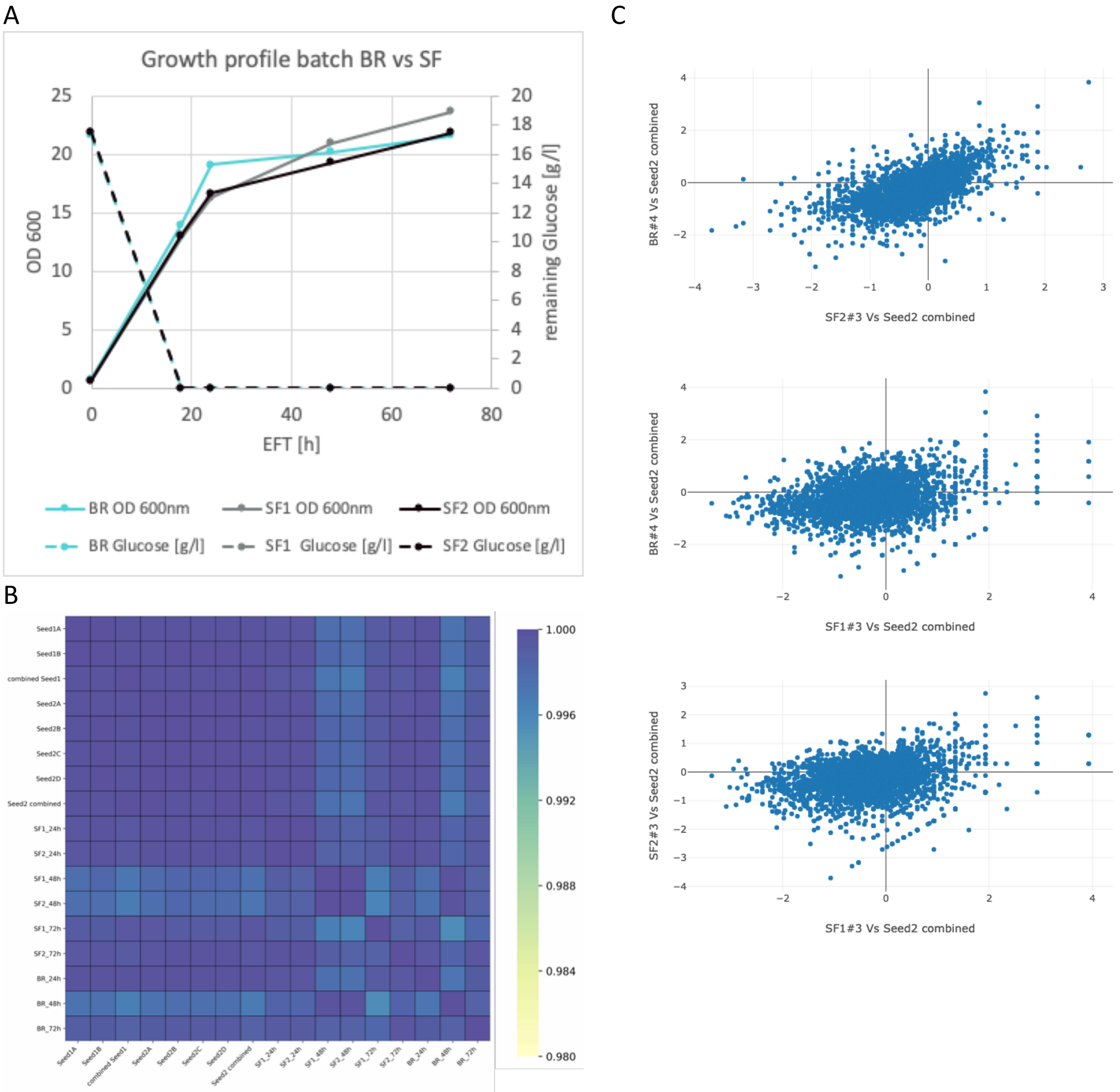


**Figure S2: Similar population dynamics in Set 1 between shake flask and batch bioreactors.** Growth (represented by $OD_{600}$) and glucose consumption (in g/L) profiles (A) for batch bioreactor (BR) and shake flasks (SF1 and SF2) during the entire cultivation time period. B)Heatmap of the Pearson correlation matrix (B) and scatter plots (C) of mutant counts for all three cultivation experiments in Set 1 growth competition experiments in comparison to seed trains.

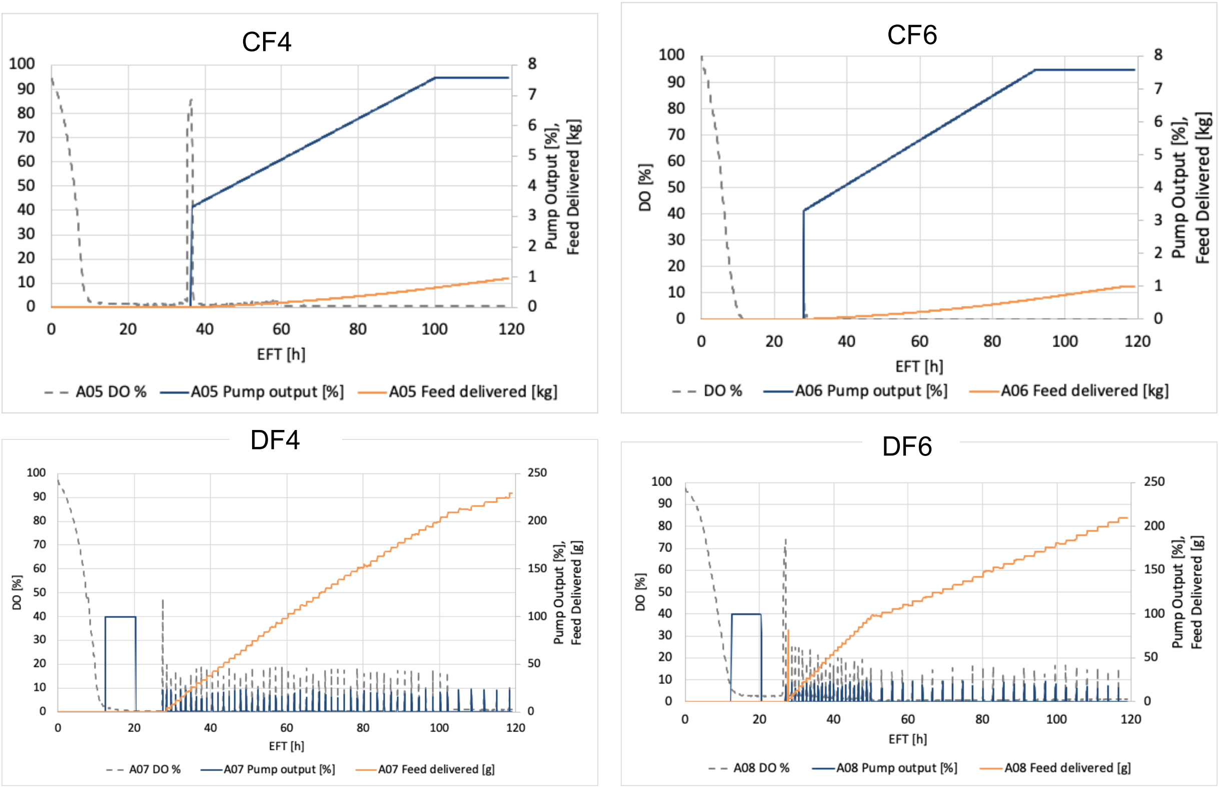


**Figure S3**: **Feeding profiles and pump output of each fed-batch bioreactor over time.** In all conditions, the feed was started upon an initial spike in dissolved oxygen (DO, grey line), indicating the consumption of available carbon sources from the batch media. For A05 and A06, a linear feed was implemented which resulted in linear addition of glucose (orange line) to the cultivation broth. For A07 and A08, the feeding pump activity (dark blue) was programmed to respond to changes in the DO (DO stat) and thus reacted to the metabolic requirements of the culture.

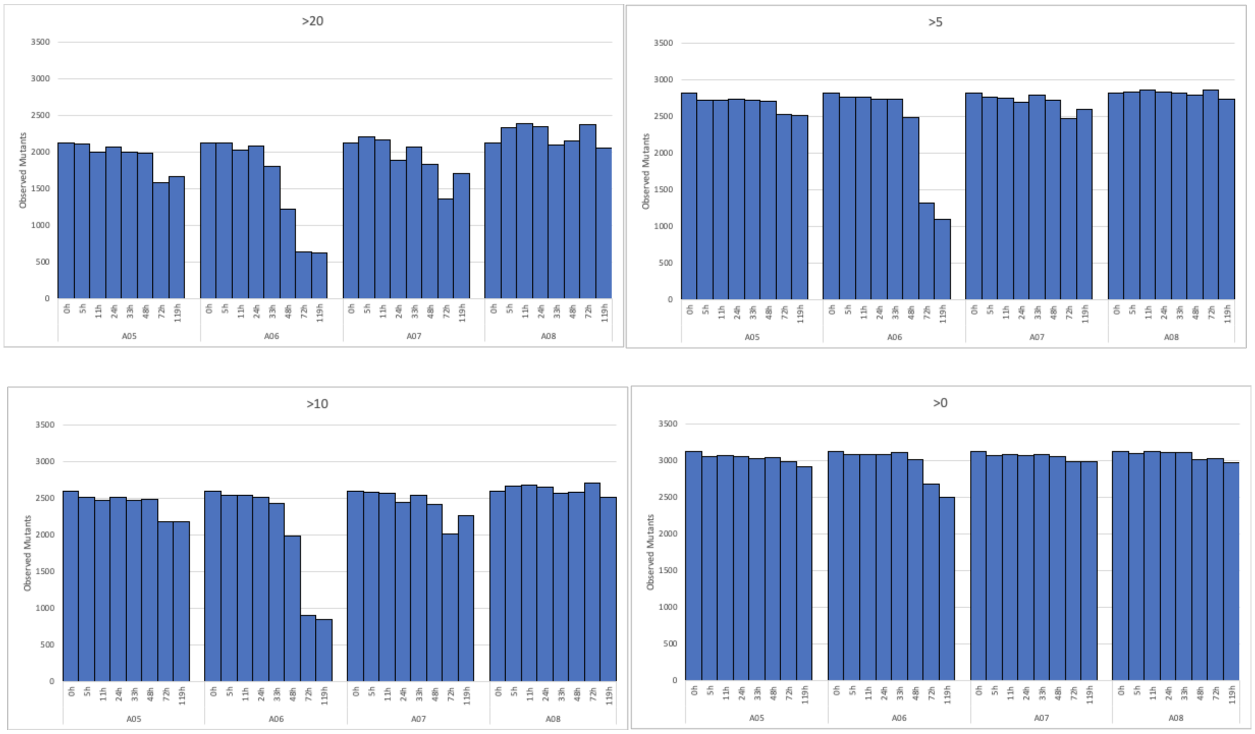


**Figure S4: Trends observed for mutants in each fed-batch bioreactor in Set 2 are independent of the mutant count threshold.** Population diversity of each fed-batch bioreactor over time using different cut off values (>20, >10, >5 and >0) in Seed 2 for the analysis. A06 had a completely different population dynamics compared to the other three bioreactors.

**A**

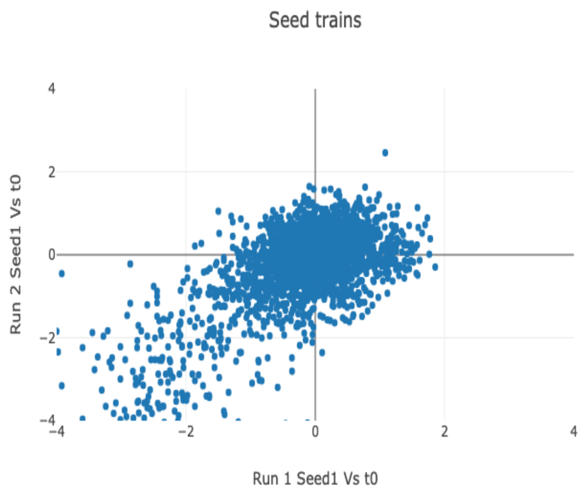


**B**

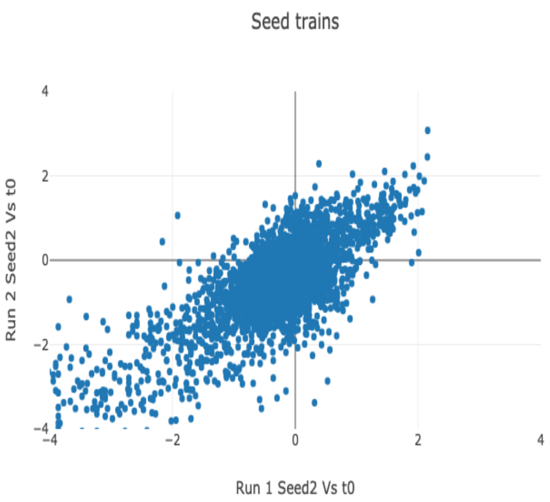


**C**

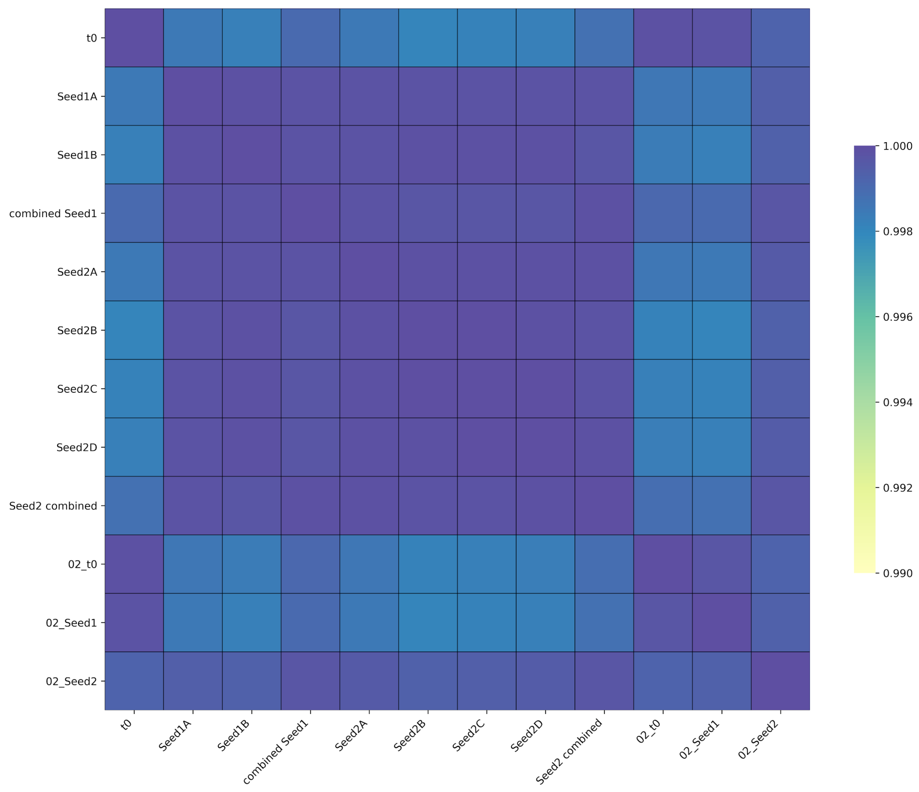


**Figure S5**: **Similar population dynamics between t0 and different seed trains.** Scatter plots (A and B) and heatmap of the Pearson correlation matrix (C) of mutant counts for t0 and individual seed trains across Set 1 and Set 2 growth competition experiments.

**Table S1: Number of strains for which fitness was measured in Set 2 and had at least 10 counts**

| -48 | -24 | 0 | Time (h) | 5 | 11 | 24 | 33 | 48 | 72 | 119 |
|---|---|---|---|---|---|---|---|---|---|---|
| t0 | Seed 1 | Seed 2 | A05 | 2425 | 2415 | 2431 | 2409 | 2391 | 2132 | 2151 |
| 2562 | 2562* | 2562 | A06 | 2440 | 2443 | 2440 | 2378 | 1976 | 896 | 844 |
| | | | A07 | 2467 | 2460 | 2392 | 2430 | 2358 | 2028 | 2204 |
| | | | A08 | 2492 | 2502 | 2513 | 2445 | 2405 | 2465 | 2372 |

*Seed 1 had 2542 mutant barcode strains with at least 10 counts.
